# Neuron-specific knockouts indicate the importance of network communication to *Drosophila* rhythmicity

**DOI:** 10.1101/639146

**Authors:** M Schlichting, MM Diaz, J Xin, M Rosbash

## Abstract

Animal circadian rhythms persist in constant darkness and are driven by intracellular transcription-translation feedback loops. Although these cellular oscillators communicate, isolated mammalian cellular clocks continue to tick away in darkness without intercellular communication. To investigate these issues in *Drosophila*, we assayed behavior as well as molecular rhythms within individual brain clock neurons while blocking communication within the ca. 150 neuron clock network. We also generated CRISPR-mediated neuron-specific circadian clock knockouts. The results point to two key clock neuron groups: loss of the clock within both regions but neither one alone has a strong behavioral phenotype in darkness; communication between these regions also contributes to circadian period determination. Under these dark conditions, the clock within one region persists without network communication. The clock within the famous PDF-expressing s-LNv neurons however was strongly dependent on network communication, likely because clock gene expression within these vulnerable sLNvs depends on neuronal firing or light.

## Introduction

Neuronal networks make myriad contributions to behavior and physiology. By definition, individual neurons within a network interact, and different networks also interact to coordinate specialized functions. For example, the visual cortex and motor output centers must coordinate to react properly to environmental changes. In a less immediate fashion, sleep centers and circadian clocks are intertwined to properly orchestrate animal physiology. The circadian system is of special interest: it not only times and coordinates physiology within neuronal tissues but also sends signals to the body to keep the entire organism in sync with the cycling external environment (Mohawk et al., 2012).

The small, circumscribed *Drosophila* clock network is ideal to address circadian communication issues. The comparable region in mammals, the suprachiasmatic nucleus, is composed of thousands of cells depending on the species. There are in contrast only 75 clock neurons per hemisphere in *Drosophila*. These different clock neurons can be divided into several subgroups according to their location within the fly brain. There are 4 lateral and 3 dorsal neuron clusters, which have different functions in controlling fly physiology (Helfrich-Förster et al., 2007).

The four small ventro-lateral neurons (sLNvs) are arguably the most important of the 75 clock neurons. This is because ablating or silencing these neurons abolishes rhythms in constant darkness (DD). They reside in the accessory medulla region of the fly brain, an important pacemaker center in many insects (Helfrich-Förster, 1997), and express the neuropeptide PDF. In addition, they are essential for predicting dawn (Depetris-Chauvin et al., 2011; Grima et al., 2004; Nitabach et al., 2002; Stoleru et al., 2004). A very recent study suggests that the sLNvs are also able to modulate the timing of the evening (E) peak of behavior (Schlichting et al., 2019). The other ventral-lateral group, the four large-ventro-lateral neurons (lLNvs), also express PDF and send projections to the medulla, the visual center of the fly brain; they are important arousal neurons (Shang et al., 2008; Sheeba et al., 2008). Consistent with the ablation experiments mentioned above, the absence of *pdf* function or reducing PDF levels via RNAi causes substantial arrhythmic behavior in DD (Renn et al., 1999; Shafer and Taghert, 2009).

Other important clock neurons include the dorso-lateral neurons (LNds), which are essential for the timing of the E peak and adjustment to long photoperiods (Grima et al., 2004; Kistenpfennig et al., 2018; Stoleru et al., 2004). Two other clock neuron groups, the lateral-posterior neurons (LPN) and a subset of the dorsal neurons (DN1s), were recently shown to connect the clock network to sleep centers in the fly central complex (Guo et al., 2018, 2016; Lamaze et al., 2018; Ni et al., 2019). The DN2 neurons are essential for temperature preference rhythms (Hamada et al., 2008), whereas no function has so far been assigned to the DN3s.

Despite these distinct functions, individual clock neuron groups are well-connected to each other. At the anatomical level, all lateral neuron clusters and even DN1 dorsal neurons send some of their projections into the accessory medulla, where they can interact. A second area of common interaction is the dorsal brain; only the lLNvs do not project there (Helfrich-Förster et al., 2007).

Several studies have investigated interactions between different clock neurons. Artificially expressing kinases within specific clock neurons causes their clocks to run fast or slow and also changes the overall free-running period of the fly, indicating that network signaling adjusts behavior (Yao and Shafer, 2014). Similarly, speeding up or slowing down individual neurons is able to differentially affect behavioral timing in standard light-dark (LD) cycles (Stoleru et al., 2005; Yao et al., 2016). A high level of neuronal plasticity within the network also exists: axons of individual cells undergo daily oscillations in their morphology (Fernández et al., 2008), and neurons change their targets depending on the environmental condition (Gorostiza et al. 2014; Chatterjee et al. 2018).

How neuronal communication influences the fly core feedback loop is not well understood. The latter consists of several interlocked transcriptional-translational feedback loops, which probably underlie rhythms in behavior and physiology (Hardin, 2011). A simplified version of the core feedback loop consists of the transcriptional activators Clock (CLK) and Cycle (CYC) and the transcriptional repressors Period (PER) and Timeless (TIM). CLK and CYC bind to E-boxes within the *period* (*per*) and *timeless* (*tim*) genes (among other clock-controlled genes) and activate their transcription. After PER and TIM synthesis in the cytoplasm, they form a heterodimer and enter the nucleus towards the end of the night. There they interact with CLK and CYC, release them from their E-box targets and thereby inhibit their own transcription. All 75 pairs of clock neurons contain this canonical circadian machinery, which undergoes daily oscillations in level. Indeed, the immunohistochemical cycling of PER and TIM within these neurons is a classic assay to visualize these molecular oscillations (Menegazzi et al., 2013).

Silencing PDF neurons stops their PER cycling, indicating an important role of neuronal firing in maintaining circadian oscillations. However, only two time points were measured, and the results were possibly confounded by developmental effects (Depetris-Chauvin et al., 2011; Nitabach et al., 2002). PDF neuron silencing also phase advances PER cycling in downstream neurons, suggesting that PDF normally serves to delay cycling in target neurons (Wu et al., 2008). This is consistent with experiments showing that PDF signaling stabilizes PER (Li et al., 2014). In addition, neuronal activation is able to mimic a light pulse and phase shift the clock due to firing-mediated TIM degradation (Guo et al., 2014).

To investigate more general features of clock neuron interactions on the circadian machinery, we silenced the majority of the fly brain clock neurons and investigated behavior and clock protein cycling within the circadian network in a standard light-dark cycle (LD) as well as in constant darkness (DD). Silencing abolished rhythmic behavior but had no effect on clock protein cycling in LD, indicating that the silencing affects circadian output but not oscillator function in a cycling light environment. Silencing similarly abolished rhythmic behavior in DD but with very different effects on clock protein cycling. Although protein cycling in the LNds was not affected by neuronal silencing in DD, the sLNvs dampened almost immediately. Interestingly, this differential effect is under transcriptional control, suggesting that some *Drosophila* clock neurons experience activity-regulated clock gene transcription. Cell-specific CRISPR/Cas9 knockouts of the core clock protein PER further suggests that network properties are critical to maintain wild-type activity-rest rhythms. Our data taken together show that clock neuron communication and firing-mediated clock gene transcription are essential for high amplitude and synchronized molecular rhythms as well as rhythmic physiology.

## Results

To investigate the effects of clock network communication on fly behavior, we silenced most adult brain clock neurons using UAS-*Kir* (Johns et al., 1999). To this end, we used the *clk856*-GAL4 driver, which is expressed in most clock neurons (Gummadova et al., 2009) and first addressed locomotor activity behavior in a 12:12 LD cycle.

Both control strains show the expected morning and evening (M and E) anticipation increases, which are normal behavioral manifestations of clock function (Fig. 1A and 1C). There is however no discernable activity anticipation in the silenced flies. Only brief activity increases are visible, precisely at the day/night and night/day transitions (Fig. 1B); these are startle responses (Rieger et al., 2003). Flies lacking PER show similar behavior (*per*^*01*^ Fig. 1D).

**Figure 1.**
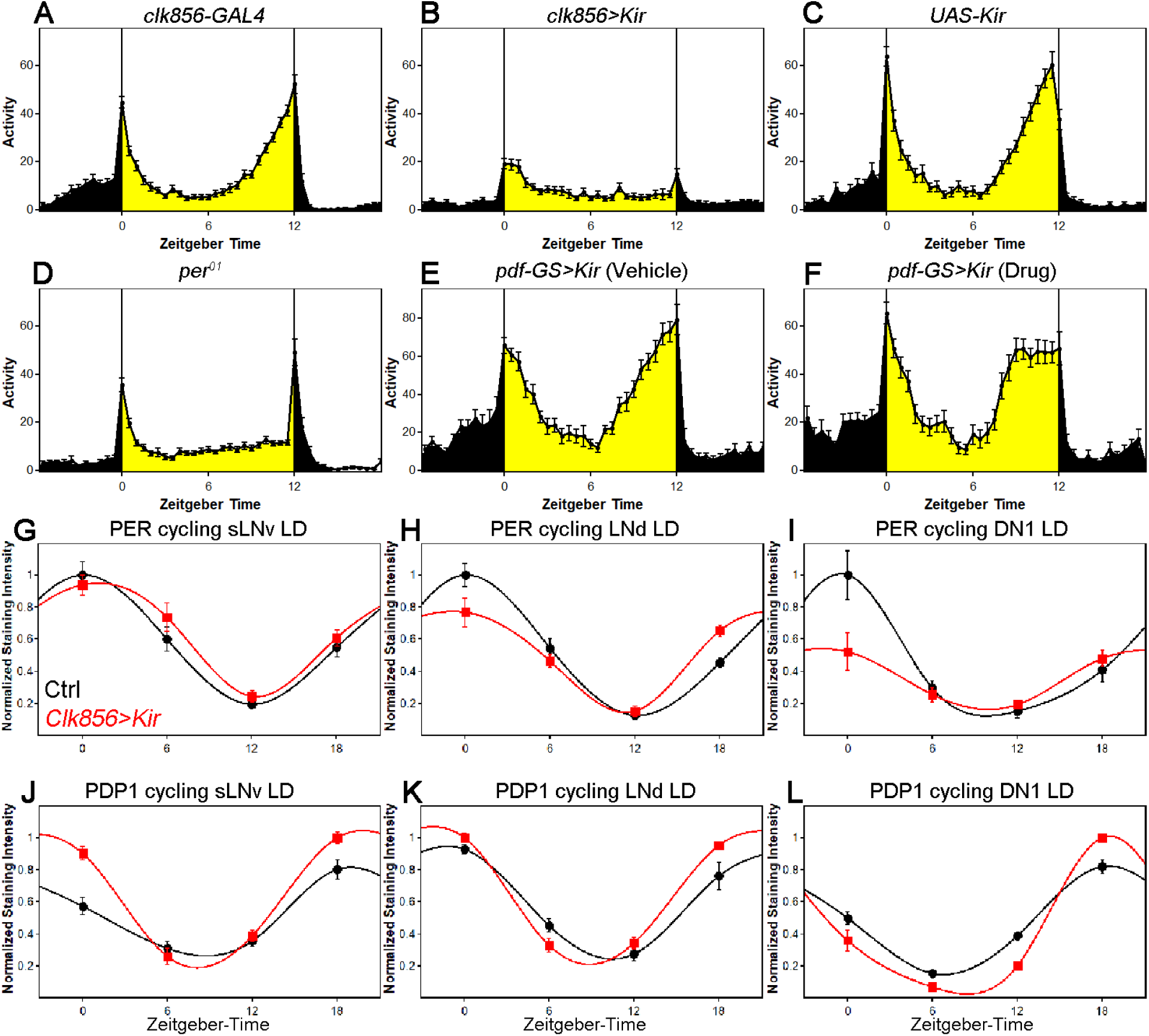
Silencing the clock network has differential effects on behavior and clock protein cycling in LD. **A-C** Silencing most of the clock network abolishes rhythmic LD behavior. GAL4 control (**A**) and UAS control (**C**) show bimodal activity patterns with M anticipation and E anticipation. Silenced flies (**B**) show no sign of anticipation neither in the morning nor in the evening. Flies show short activity increases at the transitions of day/night and night/day which are considered masking. **D** Behavior of *per*^*01*^ flies in LD 12:12. *per*^*01*^ mutants show behavior similar to *clk856>Kir* with no M and E anticipation but short reactions to the light transitions. **E-F** Silencing PDF neurons alters LD behavior. **E** *PDF-GS>Kir* on Vehicle food does not express *Kir*. Flies show the typical bimodal activity with M and E anticipation peaks. The M peak is close to lights-on (ZT0) whereas the E peak is close to lights-off (ZT12). **F** Silencing the PDF neurons by adding RU 486 to the food causes an advanced E peak, similar to *pdf*^*01*^ flies. **G-I** PER protein cycling is largely unaffected by neuronal silencing in LD. PER cycling in control brains (black data points ± SEM, pooled GAL4 and UAS) is highly synchronized with peak levels around ZT0. Silencing the clock network (red data points ± SEM) had little effect on LD PER rhythms in sLNvs (**G**) and LNds (**H**). DN1s appear dampened after silencing (**I**). **J-L** PDP1 protein cycling is largely unaffected by neuronal silencing in LD. PDP1 cycling in control brains (black data points ± SEM, pooled GAL4 and UAS) is highly synchronized with peak levels around ZT18. Silencing the clock network (red data points ± SEM) had little effect on LD PDP1 rhythms in sLNvs (**J**) and LNds (**K**) and DN1s (**L**).

To address possible developmental defects, we added *tub-GAL80ts* as an additional transgene to silence the clock network in an adult-specific manner. In this system, GAL80 is active at low temperatures (18°C) and inhibits GAL4 expression. By increasing the temperature to 30°C, GAL80 is inactivated, GAL4 is then functional and the *clk856* network silenced (McGuire et al., 2003).

At the low temperature, the controls and experimental lines show a typical wild-type bimodal activity pattern, which disappeared in experimental flies after switching to the high temperature (Suppl. Fig. 1). This shows that the *clk856>Kir* phenotype is not caused by defects during development.

We next compared the behavior to flies with silenced PDF neurons. Adult-specific silencing of the PDF neurons using the gene-switch system reduced M anticipation and significantly advanced the timing of the E peak (Fig. 1E and 1F), which reproduces previously published results (Depetris-Chauvin et al., 2011; Nitabach et al., 2002). However, the comparison with the *clk856* results shown above indicates that silencing the whole clock neuron network causes a much more severe behavioral phenotype than only silencing the PDF cells, i.e., network silencing completely abolishes rhythmic LD behavior, similar to clock mutant flies.

How does network silencing affect the circadian molecular feedback loop? To address this issue, we assayed PER as well as PDP1 protein levels in individual clock neuron clusters at four different times during the LD cycle. Both proteins show robust cycling: PER peaks at the end of the night (ZT0), and PDP1 peaks slightly earlier than PER as expected (Hardin, 2011) (Fig. 1G-1L). This indicates that network silencing has no detectable effect on clock protein timing or cycling amplitude in LD. These data further suggest that either the different neuron clocks are self-sustained, comparable to the mammalian liver, or that light can drive rhythmic gene expression even in absence of neuronal communication.

To distinguish between these possibilities, we assayed behavior and molecular cycling in constant darkness (DD). Only 17% of the silenced flies were rhythmic, indicating that network silencing causes high levels of DD arrhythmicity (Fig. 2A). To rule out developmental effects, we applied the *tub-GAL80ts* system as described above: 80 percent of the experimental flies were rhythmic at 18°C, but they were profoundly arrhythmic at 30°C with only two rhythmic flies (Suppl. Fig. 2). In contrast, adult-specific silencing of only the PDF neurons more weakly reduced rhythmicity (Fig. 2A) and also caused a short period (Fig. 2B), phenotypes that are essentially indistinguishable from those of the classical *pdf*^*01*^ mutant (Renn et al., 1999).

**Figure 2.**
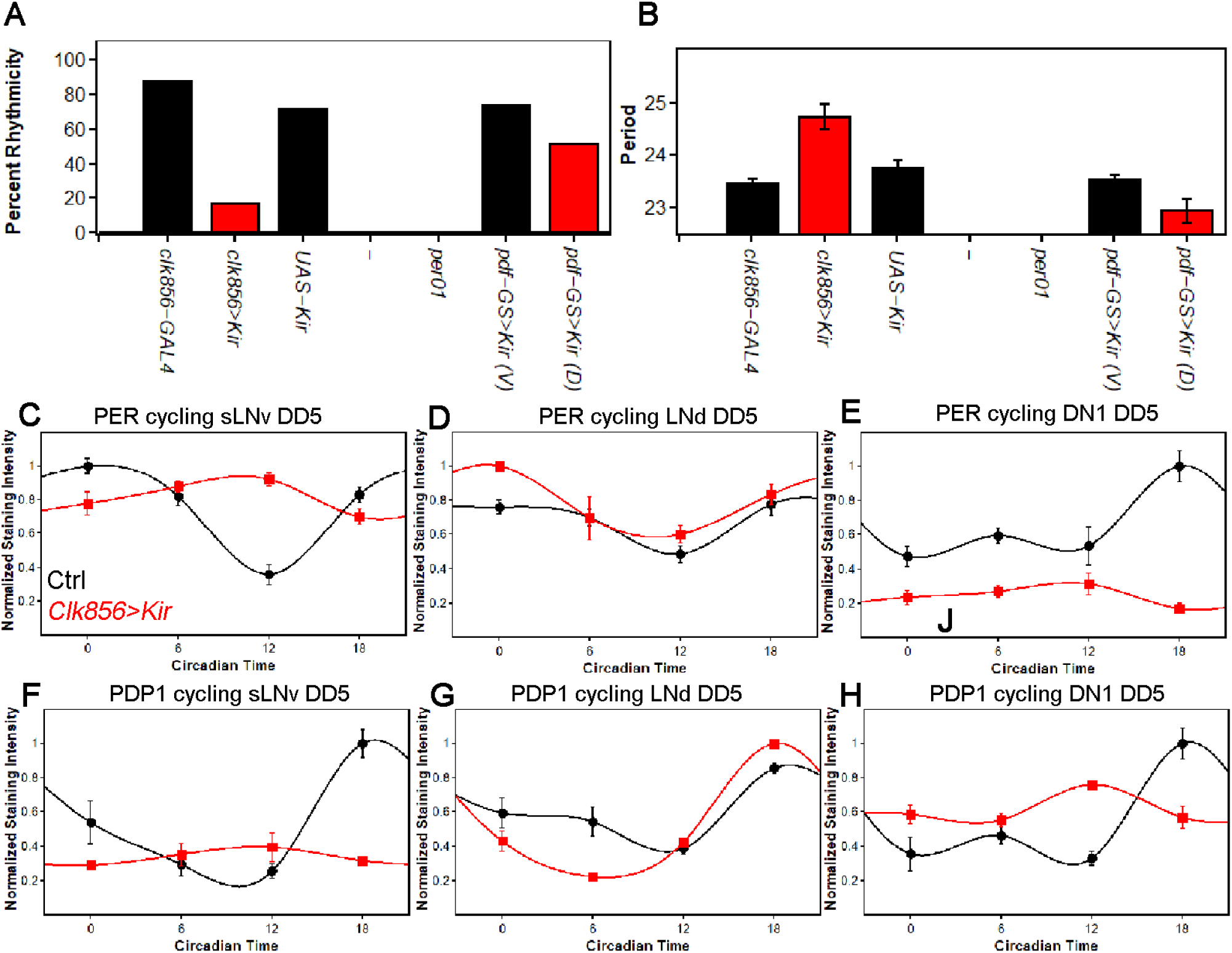
Silencing the clock network strongly affects behavior and molecular rhythms in DD **A** Percentage of rhythmic flies in DD. Silencing most of the clock neurons significantly reduces rhythmicity to less than 20 percent, suggesting that clock neuron activity is essential for rhythmic behavior output. None of the *per*^*01*^ flies were rhythmic as expected. Silencing the PDF neurons slightly decreased the level of rhythmicity. **B** Free-running period of rhythmic flies from A. The few rhythmic flies of *clk856>Kir* show a significantly longer period (F^(2,48)^=14.355 p<0.001, p<0.01 for UAS and GAL4 control). Adult-specific silencing of the PDF neurons caused a significant period shortening (p=0.0214). **C-E** PER protein cycling in DD5. PER cycling in control brains (black data points ± SEM, pooled GAL4 and UAS) is highly synchronized with peak levels around CT18. Silencing the clock network (red data points ± SEM) had variable effects on PER rhythms: The sLNvs (**C**) dampen strongly upon silencing. The LNds (**D**) show only little differences between the groups and the DN1s strongly dampen, similar to sLNvs (**E**). **F-H** PDP1 protein cycling in DD5. PER cycling in control brains (black data points ± SEM, pooled GAL4 and UAS) is highly synchronized with peak levels around CT18. Silencing the clock network (red data points ± SEM) had similar effects as observed in PER rhythms: The sLNvs (**F**) dampen strongly upon silencing. The LNds (**G**) show only little differences between the groups and the DN1s strongly dampen, but appear to have a phase-advanced PDP1 peak (**H**).

To address why network silencing has such a profound effect, we assayed PER and PDP1 protein cycling after five days in constant darkness (DD5). As expected, all assayed clock neurons from control strains maintain robust and coordinated cycling in DD (Fig. 2C-H); the sLNvs, LNds and DN1s peak slightly sooner than in LD, consistent with the slightly less than 24 hr circadian period in DD (Fig. 2B).

In striking contrast, silencing the clock network causes clock protein cycling within the individual neuronal subgroups to differ strongly from each other, in amplitude and in phase. Clock protein cycling in the LNds is least affected by neuronal silencing and with little to no change in phase or amplitude, suggesting a robust and possibly self-autonomous clock in these neurons; see Discussion (Fig. 2D and 2G). The sLNvs in contrast dampen and rapidly become arrhythmic, suggesting that these cells are rather weak oscillators and require network activity or light for proper molecular rhythms (Fig. 2C and 2F). The DN1s also dampen but less strongly. They manifest low amplitude cycling, which is phase-advanced; this intermediate situation suggests a fast and somewhat network dependent clock in DN1s (Fig. 2E and 2H). The DN2s were similar to the DN1s (data not shown). A comparable set of effects were observed in adult-specific silencing experiments (Suppl. Fig. 3).

To further address the molecular basis of the silencing dependence, we applied a fluorescent in-situ hybridization (fish) protocol to whole-mount *Drosophila* brains. Because *per* mRNA was undetectable, likely due to low expression within the clock network (data not shown and (Abruzzi et al., 2017)), we *assayed tim* mRNA cycling in LNds and in sLNvs as a proxy for clock gene transcription/mRNA levels. In control flies under LD conditions, *tim*-RNA cycles robustly in both sLNvs and LNds with a peak towards the beginning of the night as expected. In addition, clock network silencing had no effect on *tim* mRNA cycling amplitude or phase in LD, which parallels the protein cycling results (Fig. 3A and 3B). In constant darkness (DD5), the controls show robust cycling in both sLNvs and LNds as expected, but silencing causes a profound decrease in *tim* mRNA signal in the sLNvs; the LNds cycle normally (Fig. 3C and 3D). These data indicate a direct correlation between neuronal activity and *tim* RNA levels at least in the sLNvs and suggest that the silencing-mediated changes in clock protein cycling are in part transcriptional in origin.

**Figure 3.**
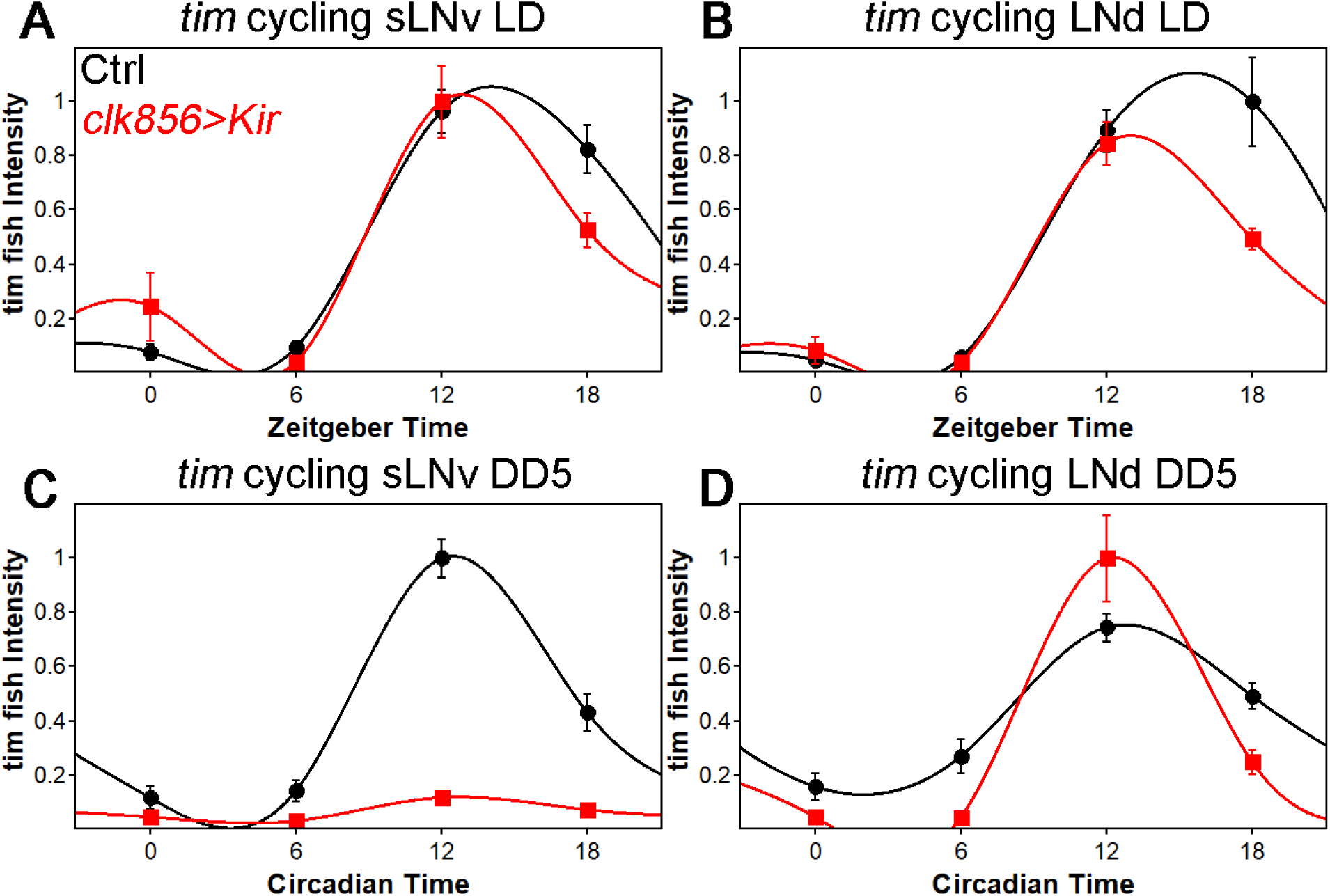
*tim* mRNA cycling in sLNvs and LNds shows similar trends as protein cycling observed by FISH. **A-B** *tim* mRNA cycling in LD 12:12 in sLNvs (**A**) and LNds (**B**). Control flies (black data points ± SEM, pooled GAL4 and UAS) show high amplitude cycling with peak levels at the beginning of the night. Silencing the clock network (red data points ± SEM) has only little effect on cycling amplitude or timing in LD. **C-D** *tim* mRNA cycling in DD5 in sLNvs (**C**) and LNds (**D**). Control flies (black data points ± SEM, pooled GAL4 and UAS) show high amplitude cycling with peak levels at the beginning of the night. Silencing the sLNvs (**C**) leads to an overall reduction of *tim* mRNA levels and a loss of rhythmicity. In the LNds (**D**) silencing did not decrease cycling amplitude or shifted peak mRNA expression.

Network silencing therefore reveals different levels of autonomy and endogenous speeds among clock neuron clusters. This leads to a drifting apart of the different subgroups from their usual well-synchronized and robust clock protein expression pattern. Interestingly, it appears that these phase differences are too big to re-establish coordinated rhythms after one week of silencing; there is no indication of rhythmic behavior upon lowering the temperature in the *tubGAL80ts* experiment (Suppl. Fig. 4).

The results to this point indicate that neuronal activity/communication is essential for rhythmicity as well as synchronized, high amplitude clock protein cycling in DD conditions. However, these results do not provide a hierarchy among the different groups, nor do they address a need for the circadian clock within these neurons. To distinguish between these possibilities and to develop a general knock-out strategy within the adult fly brain, we established a cell-specific CRISPR/Cas9 strategy to eliminate the circadian clock in individual clock neuron groups (Fig. 4A). We applied the guide protocol introduced by Port and Bullock (2016) and cloned three guides targeting the coding sequence of *per* under UAS control and generated UAS-*per-g* flies. For a first experiment, we expressed the *per-*guides and *Cas9* in most of the clock neuron network under *clk856* control and performed behavioral (Fig. 4B-4D) and immunocytochemical (Fig. 4E-4G) assays.

**Figure 4.**
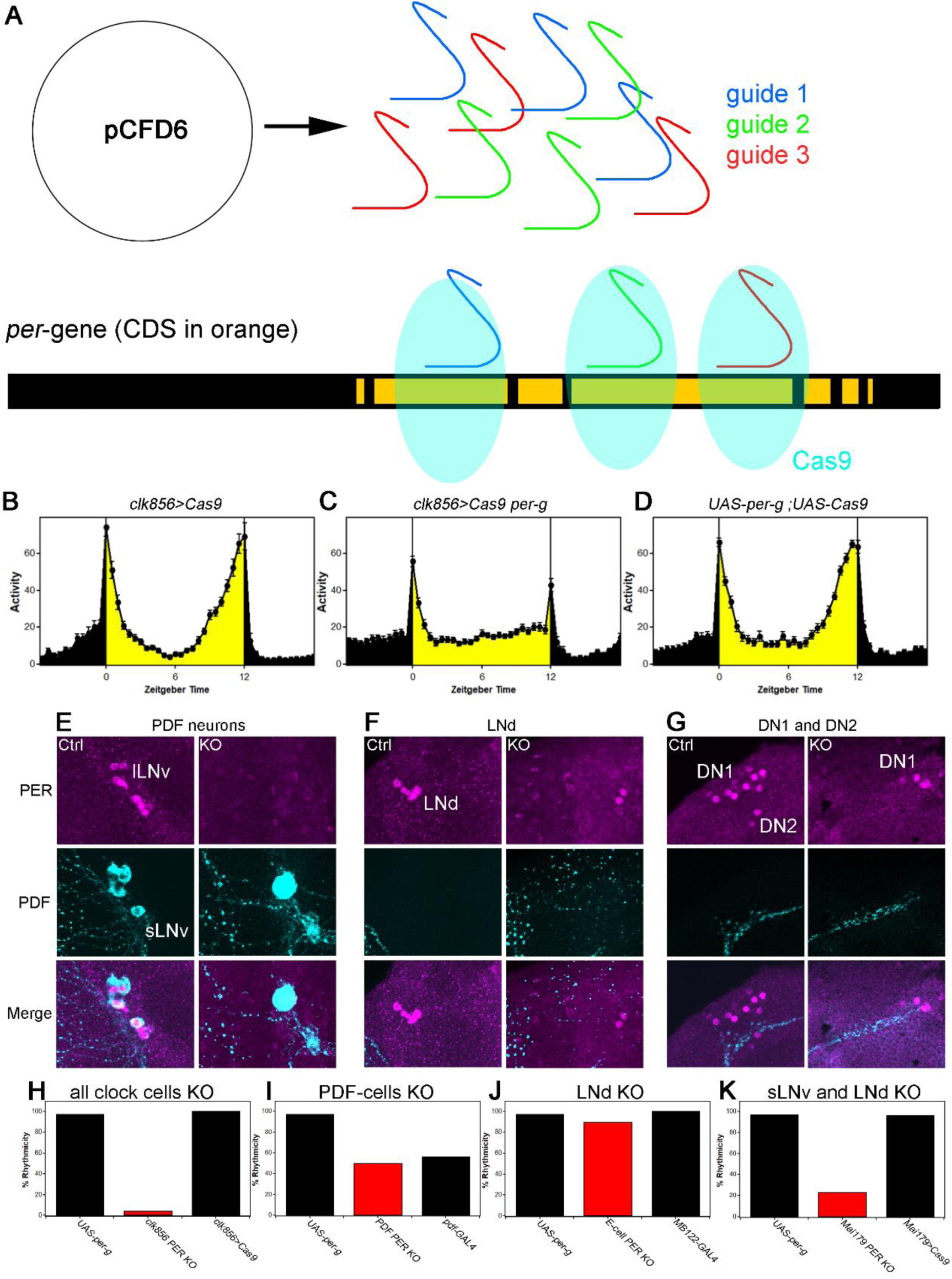
A clock in the LNds or the sLNvs can drive rhythmic behavior **A** Schematic model of cell-specific knockout (KO) strategy. We generated a UAS-*per-g* line using the pCFD6 vector, allowing us to express three guides under the control of one UAS promoter (after Port and Bullock 2016). We cloned three guides targeting the *per* CDS with guide 1 targeting the second exon shared by all transcripts and guides 2 and 3 targeting the 4th commonly shared exon. The guides will recruit the Cas9 protein and induce double-strand breaks and thereby cause mutations which lead to a non-functional protein. **B-D** Behavior of perKO using *clk856*-GAL4 in LD 12:12 reproduces *per*^*01*^ phenotype. Flies expressing *Cas9* in the majority of the clock neurons (**B**) and flies with both UAS-constructs (**D**) show bimodal activity with an M anticipation peak around lights-on and an E anticipation peak around lights-off. KO of per using *clk856*-GAL4 (**C**) abolishes M and E anticipation similar to *per*^*01*^ flies. **E-G** Immunocytochemistry of Control (*clk856>Cas9*) and KO (*clk856>Cas9, perG*) staining against PER (magenta) and PDF (cyan). Control flies show PER staining in both, sLNvs and lLNvs, whereas there is no detectable PER signal in the PDF cells in the KO strain (**E**). Similarly, we see six LNds in the control and two LNds in the experimental flies, showing that some neurons can escape (**F**) The number of PER+ DN1s is strongly reduced in the KO strain and we do not detect PER in the DN2 neurons (**G**). **H-K** A clock in LNd or PDF neurons is necessary for rhythmic behavior. *Clk856*-GAL4 mediated KO reduces rhythmicity to less than 10% (**H**). KO in PDF neurons (*pdf*-GAL4) (**I**) or in the LNds (*MB122B*-split-GAL4) (**J**) had no effect on rhythmicity. KO in both places (*Mai179*-GAL4) significantly decreases rhythmicity (**K**).

This PERKO strategy abolished M and E anticipation in LD behavior without affecting the startle responses (Fig. 4C), and it also reduced the level of DD rhythmicity to below 10%; this reproduced network silencing as well as the canonical *per*^*01*^ behavioral phenotypes (Fig. 1D and 2A). Not surprisingly perhaps given these robust phenotypes, immunohistochemistry indicates that the PERKO strategy works at more than 90% efficiency. For example, there was no detectable nuclear PER signal in all PDF cells or in the DN2s (Fig. 4E and 4G). There is also a marked reduction in the number of PER-positive DN1s in the dorsal brain; this is expected as the *clk856-*GAL4 line does not express in all DN1 neurons (Gummadova et al., 2009) (Fig. 4G). Similarly, most LNds are PER-negative. There are however two LNds that remain PER-positive for some reason (Fig. 4F), i.e., there are a few cell escapers. We note that the PERKO strategy is also effective with weaker and more narrowly expressed GAL4 lines (Suppl. Fig. 5), indicating that it can be used to investigate the contribution of clocks in individual neuron subgroups to circadian behavior.

We next addressed the contribution of clocks within individual neuron subgroups to DD rhythmicity. Previous work assigned a central role to PDF neurons and specifically to the small LNvs: ablating these cells eliminates DD rhythms, and expressing *per* in these same neurons restores DD rhythms to *per*^*01*^ flies (Grima et al., 2004; Stoleru et al., 2004). We were therefore surprised that the PERKO with *pdf*-GAL4 had no discernable effect on DD rhythmicity compared to the controls (Fig. 4I). Similarly, a PERKO in the cells important for controlling E activity (E cells: 3 LNds and the 5th sLNv) with *MB122B*-split-GAL4 had no effect on rhythmicity (Fig. 4J). However, a PERKO in both groups achieved with *Mai179*-GAL4, lowered rhythmicity to less than 20% (Fig. 4K). Similar results were obtained with *dvPDF*-GAL4, which expresses in similar neuron groups (data not shown).

To address whether other neurons have similar effects, we expressed the PER guides elsewhere: knockout in the retina (*GMR*-GAL4), glial cells (*repo*-GAL4) or DN1s (*clk4.1M*-GAL4 and *AstC*-GAL4) did not affect rhythmicity (Suppl. Fig. 6). These findings taken together suggest that a clock in either of two key places, the sLNvs or the LNds, can drive rhythmic behavior.

We also assayed the free-running DD periods of flies lacking PER in individual neuron subgroups (Fig. 5A). These periods did not change if the PERKO was in the dorsal brain and/or in the large PDF neurons, the lLNvs. However, a PERKO in the E cells and with drivers expressing in these cells plus some dorsal neurons results in a slight but significant period lengthening of approximately 0.5hr. In contrast, a PERKO in most of the lateral neuron clusters gave rise to a short period. These two sets of period phenotypes taken together suggest that the clocks in the two different key neuron subgroups collaborate to achieve the intermediate and close to 24h period characteristic of wild-type flies.

**Figure 5.**
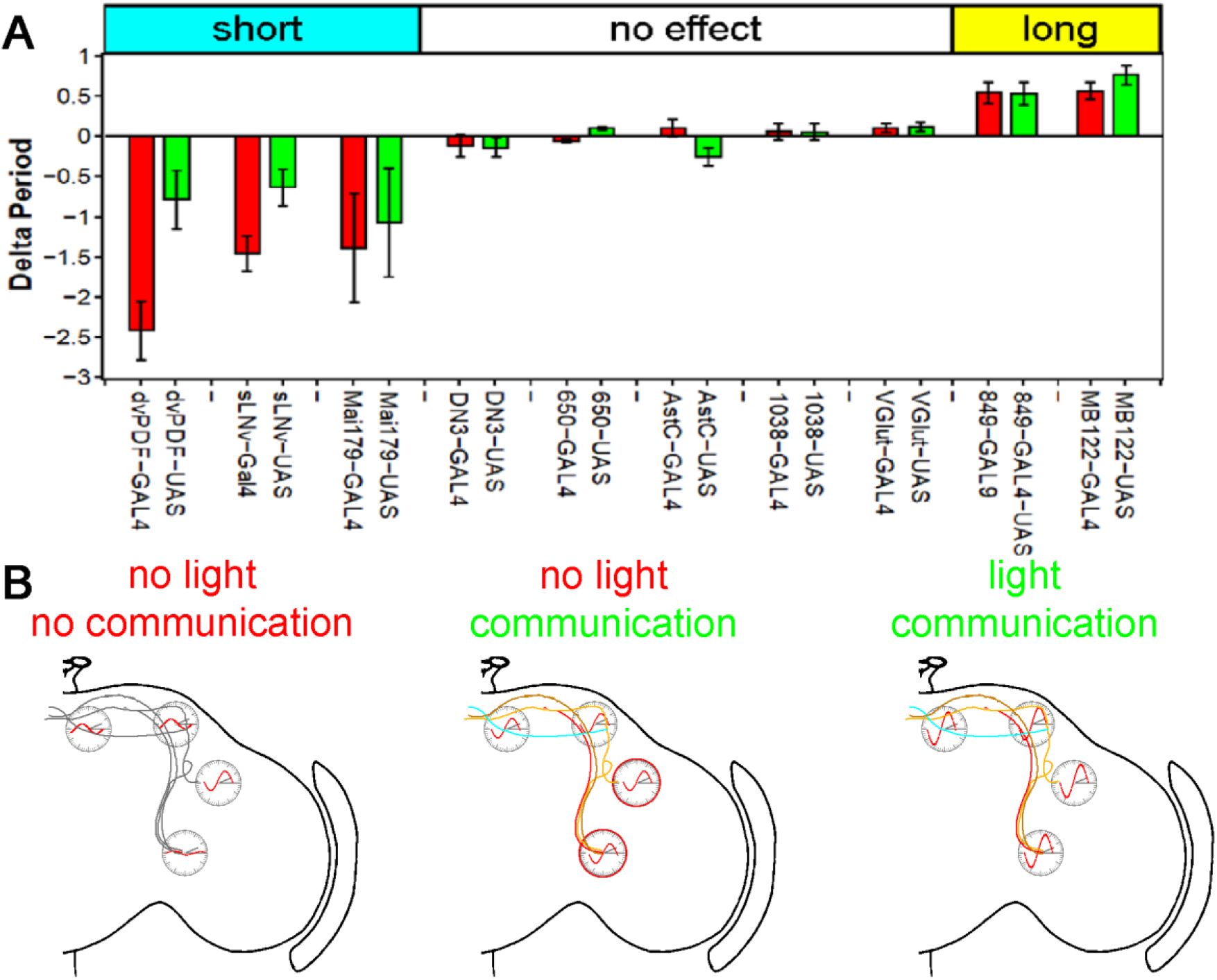
KO of PER in subsets of neurons changes free-running period. **A** Changes of free-running period upon KO. Red bars represent the change of period between the KO line and the GAL4 control (± SEM), green bars represent the change of period between the KO line and the UAS control (± SEM). KO in most of the lateral neurons (*Mai179*-GAL4, *dvPDF*-GAL4) causes a shift towards short periods. KO in the dorsal brain (*clk4.1M*-GAL4, *AstC*-GAL4, *GMR-ss00657*-GAL4, *GMR-ss00650*-GAL4, *GMR-ss01038*-GAL4, *VGlut*-GAL4) does not affect the period, whereas KO in the LNds (*MB122B*-GAL4, *GMR-ss00849*-GAL4) lengthens the period. **B** Model of neuronal communication and light influencing the Drosophila clock machinery. Silencing the clock network in DD causes a damping of molecular oscillations and a drifting apart from the common phase as indicated by the red waves. If network communication is allowed, the different neuronal sub-clusters are mostly in sync and show robust cycling, suggesting neuronal communication is essential for molecular oscillations. In a normal LD cycle light drives high amplitude and synchronized cycling even in the absence of neuronal communication, establishing a hierarchy of synchronization cues with light on the top.

## Discussion

The central clock of animals is essential for dictating the myriad diurnal changes in physiology and behavior. Knocking out core clock components such as *period* or *Clock* severely disrupts circadian behavior as well as molecular clock properties in flies and mammals (Allada et al., 1998; Gekakis, 1998; Konopka and Benzer, 1971). Here we show that similar behavioral effects occur when we silence the central clock neurons and thereby abolish communication within this network and with downstream targets, i.e., fly behavior becomes arrhythmic in LD as well as DD conditions and resembles the phenotypes of core clock mutant strains (Konopka and Benzer, 1971).

Despite the loss of all rhythmic behavior, silencing did not impact the molecular machinery in LD conditions: PER and PDP1 protein cycling was normal. These findings suggest that 1) rhythmic behavior requires clock neuron output, which is uncoupled from the circadian molecular machinery by network silencing, and 2) synchronized molecular rhythms of clock neurons do not require neuronal activity. These findings are in agreement with previous work showing that silencing the PDF neurons had no effect on PER cycling within these neurons (Depetris-Chauvin et al., 2011; Nitabach et al., 2002; Wu et al., 2008). The results presumably reflect the strong effect of the external light-dark cycle on these oscillators.

In DD however, the individual neurons change dramatically: the different neurons desynchronize, and their protein cycling damps to different extents. Interestingly, sLNv cycling relies most strongly on neuronal communication: these neurons cycle robustly in controls but apparently not at all in the silenced state. sLNvs were previously shown to be essential for DD rhythms (Grima et al., 2004; Stoleru et al., 2004). Unfortunately, the sensitivity of immunohistochemistry precludes determining whether the molecular clock has actually stopped or whether silencing has only (dramatically) reduced cycling amplitude. However, a simple interpretation of the adult-specific silencing experiment favors a stopped clock: decreasing the temperature to 18 degrees after a week at high temperature failed to rescue rhythmic behavior. A similar experiment in mammals gave rise to the opposite result, suggesting an effect of firing on circadian amplitude in that case (Yamaguchi et al., 2003). However, we cannot at this point exclude a different explanation, for example a too large phase difference between the different neuronal subgroups to reverse after a week without communication.

In either case, a stopped clock or an effect on clock protein oscillation amplitude, these results make another link to the mammalian literature: modeling of the clock network suggests that different neurons resynchronize more easily if the most highly-connected cells are intrinsically weak oscillators (Webb et al., 2012). The sLNvs are essential for DD rhythms, known to communicate with other clock neurons (Grima et al., 2004; Stoleru et al., 2004) and are situated in the accessory medulla; this is an area of extensive neuronal interactions in many insects (Reischig and Stengl, 2003). These considerations rationalize weak sLNv oscillators.

An important role of interneuron communication in DD is in agreement with previous work showing that altering the speed of individual neuron groups can change the phase of downstream target neurons (Yao and Shafer, 2014). An important signaling molecule is the neuropeptide PDF: its absence changes the phase of downstream target neurons, and silencing PDF neurons causes an essentially identical phenotype to the lack of PDF (Im et al., 2011; Lin et al., 2004; Wu et al., 2008). However, the effects reported here are much stronger and show different levels of autonomy than PDF ablation, suggesting that other signaling molecules and/or the neuronal activity of additional clock neurons are essential to maintain proper rhythmic clock protein expression.

To address these possibilities, we took two approaches. First, we investigated clock gene RNA levels after silencing. The goal was to assess whether the damping of silenced neurons is under gene expression control, likely transcriptional control. Indeed, *tim* mRNA profiles nicely reproduced the protein cycling profiles: robust cycling of all (assayed) clock neurons was maintained in LD even with silencing, but *tim*-mRNA levels in the sLNvs stopped cycling in DD; in contrast, robust cycling was maintained in the LNds (Fig 2D and 2G). This suggests that the changes in protein cycling amplitude and also possibly phase are under transcriptional control. Importantly, the *tim* signal in the sLNvs disappeared upon silencing, suggesting that neuronal activity promotes clock gene transcription at least in this subset of neurons. This recapitulates for the first time in *Drosophila* the robust positive relationship between neuronal firing and clock gene transcription in mammals (Shigeyoshi et al., 1997). To date, *Drosophila* neuronal firing had only been connected to post-transcriptional clock protein regulation, namely TIM degradation (Guo et al., 2014). Conceivably, these two effects are connected: TIM degradation might be required to relieve transcriptional repression and maintain cycling.

The second approach was a cell-specific knockout strategy, applied to the clock neuron network. We generated three guides targeting the CDS of *per* and also expressed CAS9 in a cell-specific manner. The guides caused double strand breaks in the *per* gene, which in turn led to cell-specific *per* mutations. This adult brain knockout strategy worked reliably and specifically, in glial cells as well as neurons, with a greater than 90% efficiency and with no apparent background effects (Fig. 4B-G). We have successfully used this strategy to knock out most if not all *Drosophila* GPCRs (data not shown) and believe it will be superior to RNAi for most purposes. Importantly, expression of the guides with the *clk856*-GAL4 driver phenocopied *per*^*01*^ behavior (Fig. 4C and 4H). To focus on individual clock neurons, we generated cell-specific knockouts in different clock neurons. To our surprise, a PERKO in the PDF cells did not increase the level of arrhythmicity. Only a PERKO in most lateral neurons, E cells as well as PDF cells, generated high levels of arrhythmic behavior. As PDF cell ablation also causes high levels of arrhythmicity (Stoleru et al., 2004), the data shown here suggest that the LNds can drive the rhythmic output of key sLNv genes even in the absence of a clock in these neurons. Consistent with this interpretation, recent data indicate that the LNds project dendritic as well as axonal arborizations into the accessory medulla, the location of the sLNvs, indicating extensive communication between these two important subgroups of clock neurons (Schlichting et al., n.d.). This interpretation is further supported by examining the free-running period of the cell-specific knockouts: Ablating *per* in the LNds causes a long period, whereas ablating PER in most lateral clock neurons causes a short period phenotype. These data suggest that interactions between the sLNvs and other clock cells, perhaps within the accessory medulla, are essential for the close to 24h speed of the overall brain and behavioral clock.

While this manuscript was being written, we became aware of two other studies addressing the contribution of different circadian subgroups and neuronal interactions to *Drosophila* rhythms. The strategy, results and conclusions in the first study overlap extensively to what we report here (Delventhal et al., cosubmitted paper). That work exploited guides against *tim* as well as against *per* and thoroughly characterized the efficacy of the cell-specific knockout strategy, including effects on mRNA cycling.

The second study is recently published and similarly highlights the dependence of DD rhythms on network properties (Bulthuis et al., 2019). However, they report a decrease in rhythmicity upon knockout of the circadian clock in PDF cells, an effect that neither we nor Delventhal observed. This difference may be due to their knockout strategy, namely, overexpression of a dominant negative Cycle isoform within PDF cells. This protein may have effects on gene expression beyond knocking out the circadian clock. This further suggests that the conceptually simpler PERKO strategy has fewer side effects and is therefore superior.

Some of the communication properties described here resemble what has been found in mammalian systems. For example, decreasing neuronal interactions by creating sparse SCN cultures changes the free-running period and activity phase of individual neurons (Welsh et al., 1995). This suggests that communication is also critical for circadian phase and period determination in mammals. Nonetheless, fly clock cells may be even less cell-autonomous than what has been described for mammals (reviewed in (Evans, 2016)). First, the fly system may be particularly dependent on light. For example, peripheral fly clocks appear strongly light-dependent in contrast to what has been described for mammalian liver (reviewed in (Ito and Tomioka, 2016). Although much of the fly data could reflect cellular asynchrony in constant darkness, circadian cycling in the periphery crashes rapidly under these conditions and resembles the strong and rapid non-cycling that occurs in the sLNvs upon silencing in DD. Notably, fly cryptochrome but not mammalian cryptochrome is light-sensitive (reviewed in (Michael et al. 2017) and probably contributes to the light-dependence of fly peripheral clocks. This is also because light can directly penetrate the thin insect cuticle, which probably contributes to making the fly brain less dependent on ocular photoreception than the mammalian brain. However, some fly clock neurons do not express Cryptochrome, suggesting that the fly clock system is dependent on network interactions even in a light-dark cycle (Benito et al., 2008; Yoshii et al., 2008). These considerations suggest that the fly circadian network is an attractive object of study not only because of its limited size of 75 neuron pairs but also because of its strong dependence on neuronal communication.

## Material and Methods

### Fly strains and rearing

The following fly lines were used: *clk856-*GAL4 (Gummadova et al., 2009), UAS-*Kir2.1* (BL: 6595), *pdf-GS*-GAL4 (Depetris-Chauvin et al., 2011), *per*^*01*^ *(Konopka and Benzer, 1971), mai179-*GAL4 (Grima et al., 2004), *MB122B-*GAL4 (Guo et al., 2017), *pdfM*-GAL4 (Renn et al., 1999), *clk4.1M*-GAL4 (Zhang et al., 2010), UAS-*Cas9.P2* (BL 58986), AstC-GAL4 (BL: 52017), dvPDF-GAL4 (Guo et al., 2014), *w;CyO/Sco;MKRS/TM6B* (BL: 3703), VGlut-GAL4 (BL: 60312), GMR-ss0650-GAL4, GMR-ss01038-GAL4, GMR-ss00849-GAL4, GMR-ss00367-GAL4, GMR-ss00681 (Liang et al., 2019), GMR-GAL4 (BL: 1104), repo-GAL4 (BL: 7415), tub-GAL80ts (BL: 7018). The SS00849, SS00367, SS01038, SS00645, SS00650 lines were made and characterized by H. Dionne and A. Nern in the laboratory of G. Rubin (Janelia
Research Campus). All flies were reared on standard cornmeal medium at a temperature of 25°C, with the exception of adult-specific silencing experiments for which flies were raised at 18°C.

### Fly line generation

We generated a UAS-*per-g* line following the protocol published by (Port and Bullock, 2016). In short, we digested the pCFD6 Vector (addgene #73915) with BbsI, PCR amplified two PCR fragments carrying three guides targeting the CDS of *per* and performed a Gibson Assembly to include those in the pCFD6 backbone. Positive clones were sent for injection to Rainbow Transgenic Flies Inc (Camarillo, CA, USA) and the transgene was inserted into the second chromosome by phi-recombinase using BL 8621. Flies were crossed to *w*^*1118*^ for screening and positive individuals were balanced using BL 3703. The following guide sequences were used:

*per* guide1: GGCAGAGCCACAACGACCTC

*per* guide2: CAAGATCATGGAGCACCCGG

*per* guide3: GAGCAAGATCATGGAGCACC

### Behavior recording and data analysis

Individual 2-6 days old male flies were singularly transferred into glass tubes (diameter 0.5mm) with food (2% agar and 4% sucrose) on one end and a cotton plug to close the tube on the other end. The tubes were placed into *Drosophila* Activity Monitors (DAM, Trikinetics) in a way that the infrared light beam was located in the center of the tube. A computer measured the number of light-beam interruptions caused by the movement of the fly in one-minute intervals. We recorded the behavior of all flies at a constant temperature of 25°C for 5-7 days under standard light-dark conditions of 12h light and 12h darkness (LD12:12) followed by constant darkness (DD) for at least 6 days. For performing adult-specific silencing experiments, we raised the flies at 18 degrees and performed 2 separate sets of experiments: In the LD experiment, we recorded the behavior of the flies for 3 days at 18°C and switched to 30°C to silence the neurons and follow the behavioral change within the same sets of flies. In a second set of experiments, we raised the two groups of flies at 18°C and then performed LD to DD experiments either at 18°Cor 30°C. In the 30°C experiment, we decreased the temperature back to 18°C at Circadian Time (CT) 0 after 6 days in DD to investigate possible emergence of rhythmic behavior after silencing. We continued recording the behavior for 6 more days in DD at 18°C.

We generated actograms using ActogramJ (Schmid et al., 2011). We next generated average activity profiles of at least the last 3 days of LD condition as previously described (Schlichting and Helfrich-Förster, 2015). Each experiment consists of at least 2 biological repeats. DD analysis was performed using chi2-analysis. Statistical analysis was performed using a student’s t-test or one-way ANOVA followed by post-hoc Tukey analysis.

### Immunohistochemistry

2-6 days old male flies were entrained in LD 12:12 at 25°C for three days and collected at ZT0 to analyze the CRISPR/Cas9 knockout strategy. To investigate clock protein cycling, 2-6 days old male flies were entrained in LD 12:12 at 25°C for 5 days and collected in 6h intervals around the clock. Similarly, flies were entrained for 5 days and released into DD for 5 more days to obtain cycling data at DD5.

The whole flies were fixed for 2h 45min in 4% paraformaldehyde (PFA) in phosphate-buffered saline (pH=7.4) including 0.5% TritonX (PBST). The flies were rinsed 5 times for 10 min each with PBST and subsequently the brains were dissected in PBST. Brains were blocked in 5% normal goat serum (NGS) in PBST for 3h at room temperature (RT). The primary antibody (rabbit anti-PER, 1:1000, (Stanewsky et al., 1998), mouse anti-PDF, 1:500, *Drosophila* Studies Hybridoma Library (DSHB), C7 and guinea-pig anti PDP1, 1:2000, (Benito et al., 2007)) was applied overnight at RT and the brains subsequently rinsed 5 × 10mins with PBST. Secondary antibodies (Alexa, Fisher Scientific, 1:200) were applied for 3h at RT. Afterwards, the brains were rinsed 5 × 10 mins with PBST and mounted on glass slides using Vectashield (Vector Laboratories INC., Burlingame, CA, USA) mounting medium.

Confocal microscopy was performed using a Leica SP5 microscope. Sections of 1.5 um thickness were obtained. Laser settings were kept constant across genotypes to obtain comparable results. Image acquisition was performed using Fiji. Staining intensity was assessed by quantifying the brightest 3×3 pixel area of individual neurons of at least 5 brains per timepoint. Each experiment consists of at least 2 biological repeats. Three different background intensities were determined the same way and subtracted from the neuronal intensity. Data points represent average and SEM.

### Fluorescent in-situ hybridization (fish)

2-6 days old male flies were entrained for 5 days in LD 12:12 and collected in 6h intervals around the clock. In a second set of experiments, flies were released into DD for 5 days and collected in 6 h intervals. Flies were dissected fresh under red light to avoid phase-shifting the molecular machinery. Brains were subsequently fixed in 4% PFA in PBS for 55 min at RT. Afterwards, brains were washed 3 × 10min in PBST and dehydrated as described in(Long et al., 2017). Brains were kept in 100% EtOH until all time points were collected and all further steps were done simultaneously as described in (Long et al., 2017).

A set of 20-probe sequences were designed for the entire *pdf* mRNA sequence and conjugated with Quasar 570 (Stellaris Probes, Biosearch Technologies, CA, USA). The *tim* probes consist of a set of 48-probe sequences against the entire *tim* mRNA sequence, including the 5’ and 3’ untranslated regions. The *tim* probes were conjugated with Quasar 670 dye (Stellaris Probes, Biosearch Technologies, CA, USA). Probes were diluted to a stock concentration of 25 μM and aliquoted in −20 °C. The final concentration of *pdf* probes and *tim* probes were 250 nM and 750 nM, respectively.

Brains were mounted on glass slides using Vectashield mounting medium (Vector Laboratories INC., Burlingame, CA, USA) and scanned using a Leica SP5 microscope in 1.5 um sections. All samples were scanned in one session to avoid signal loss. Fluorescence intensity was assessed by quantifying the brightest 3×3 pixel area of individual neurons of at least 5 brains. Each experiment consists of at least 2 biological repeats. Three different background intensities were determined the same way and subtracted from the neuronal intensity. Data points represent average and SEM.

## Acknowledgements

We would like to thank Dr. Fang Guo and Dr. Katharine Abruzzi for discussions and comments on the manuscript. Also, we are thankful to Dr. Norbert Perrimon and Dr. Paul Hardin for providing fly lines and antibodies. We thank H. Dionne, A. Nern and G. Rubin (Janelia Research Campus) for providing the unpublished split-GAL4 lines: SS00849, SS00367, SS01038, SS00645 and SS00650. Stocks obtained from the Bloomington *Drosophila* Stock Center (NIH P40OD018537) were used in this study. This work was supported by the Howard Hughes Medical Institute (HHMI). M.S. was sponsored by a DFG research fellowship (SCHL2135 1/1).

---

**Supplement Figure 1.**
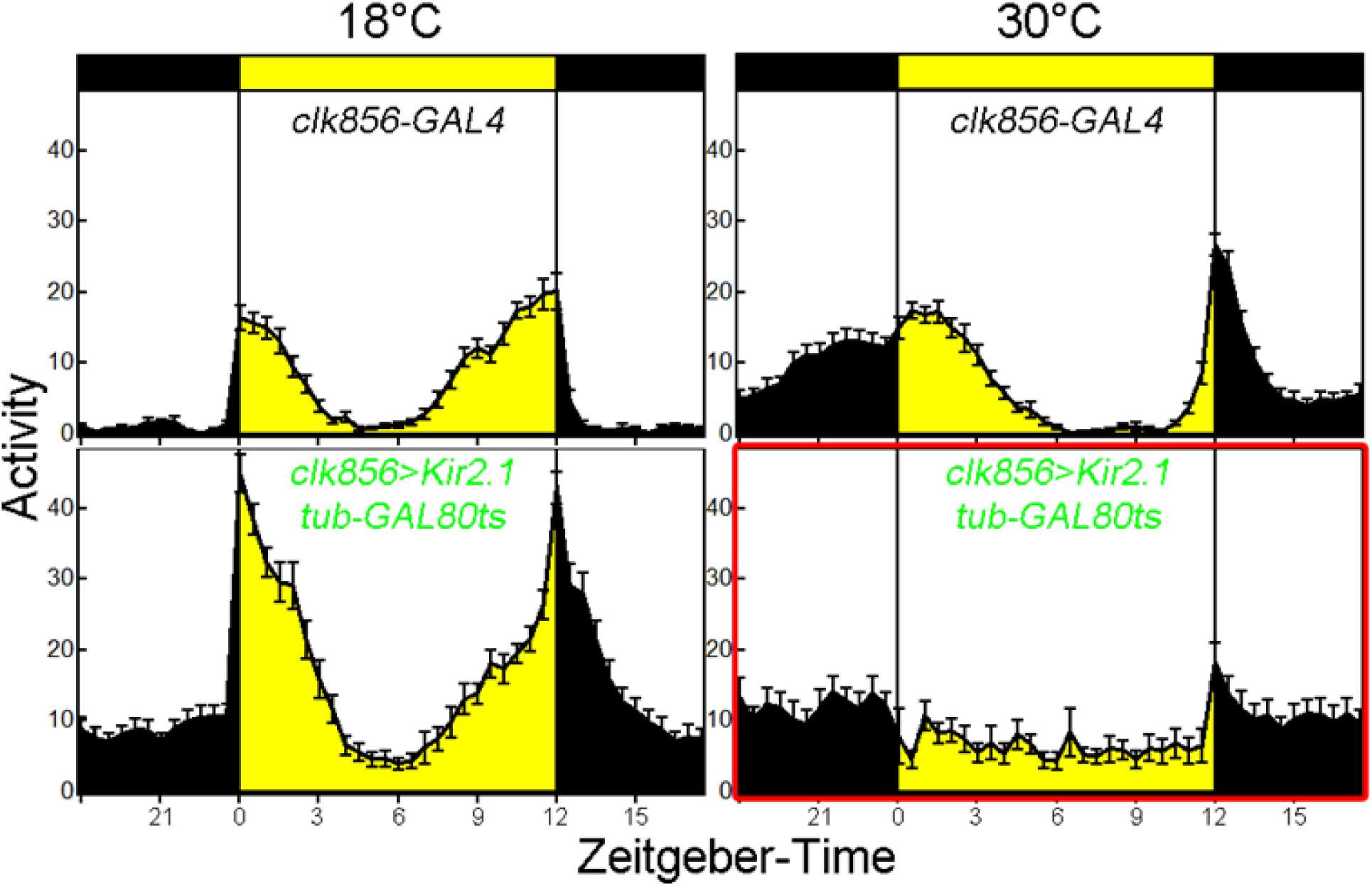
Adult-specific silencing of most clock neurons causes arrhythmic LD behavior. Control (*clk856*-GAL4) and experimental (*clk856-*GAL4 UAS*-Kir tub-GAL80ts*) flies show bimodal activity at 18 degrees with an M and an E peak of activity. M anticipation is reduced in both cases due to the low temperature. At 30 degrees, control flies (upper panel) show a bimodal activity pattern with an M anticipation peak and an E anticipation peak. The E peak is delayed due to high temperatures. The experimental flies (lower panel) show no M or E peak at 30 degrees.

**Supplement Figure 2.**
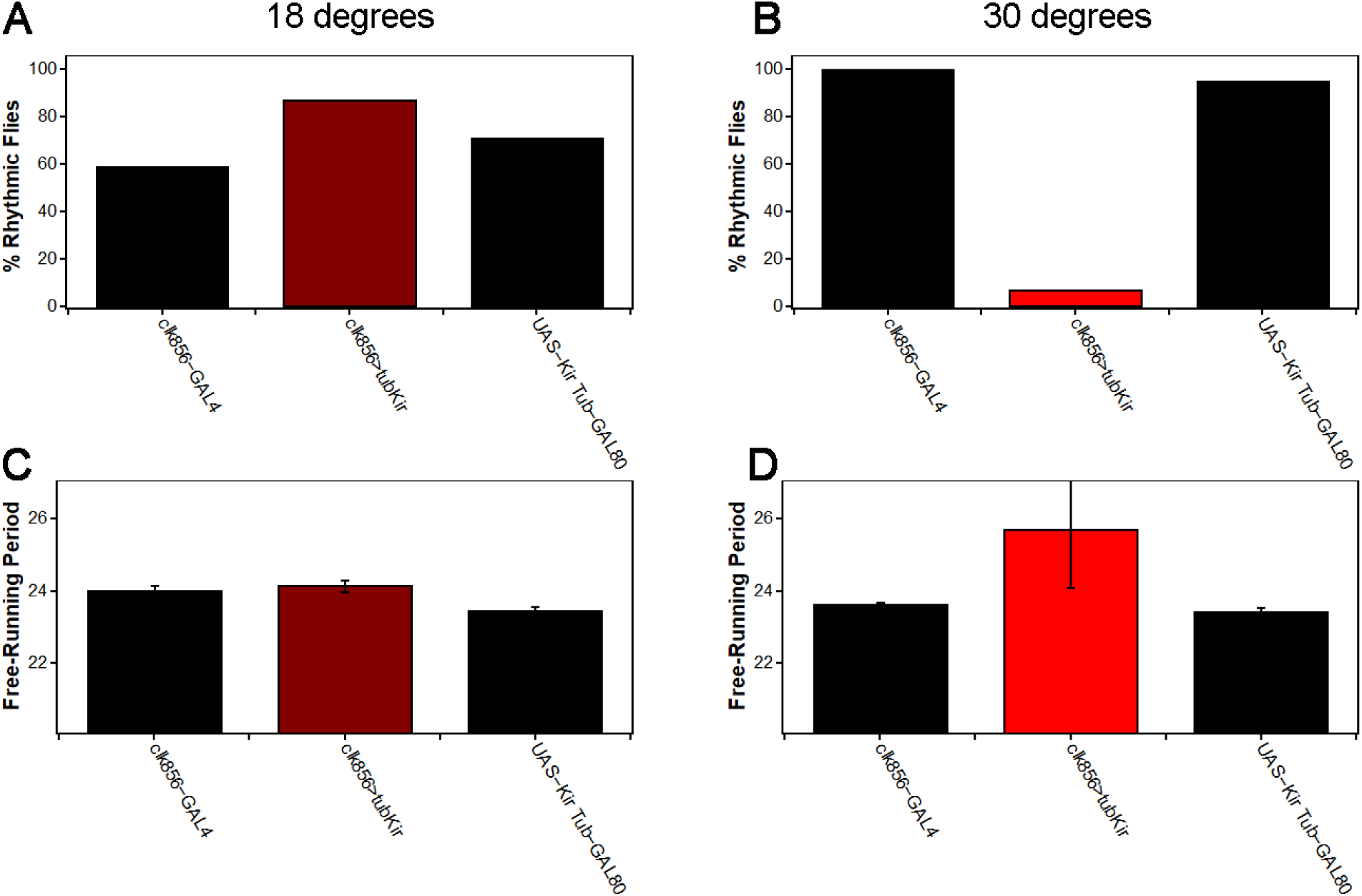
Adult-specific silencing of most clock neurons causes arrhythmic DD behavior. **A** At 18 degrees we observe no decrease in rhythmicity in *clk856>tubKir* (dark red) compared to control (black) flies as expected. Increasing the temperature to 30 degrees (**B**) activates GAL4 and hence silences the neurons. *Clk856>tubKir* (red) flies get arrhythmic, whereas both controls (black) show high levels of rhythmicity. We observed no effect on free-running period at 18 degrees (**C**) but flies experimental flies showed the tendency towards a long period at 30 degrees (**D**), similar to *clk856>Kir* (Fig. 2B).

**Supplement Figure 3.**
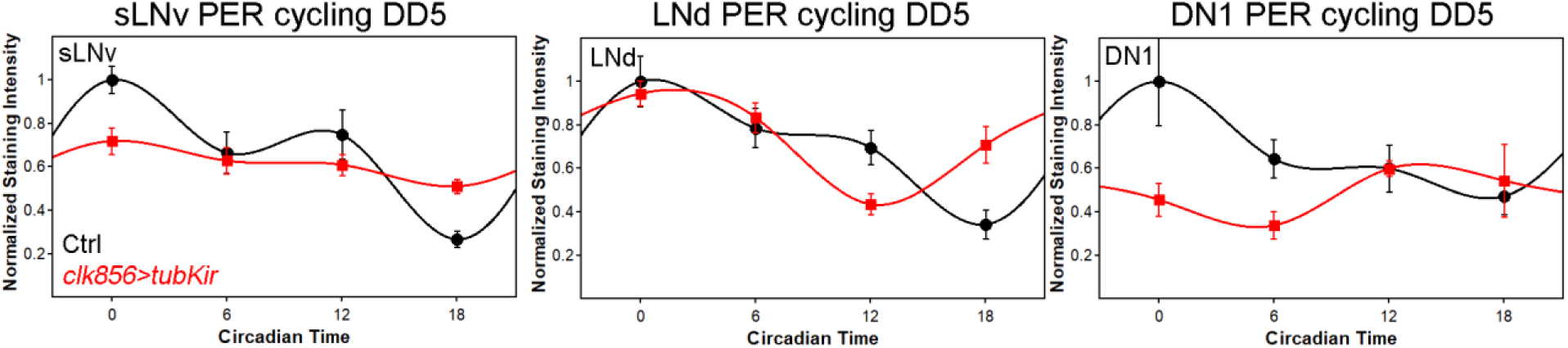
Adult-specific silencing reproduces PER cycling profiles. Flies were raised at 18 degrees and entrained at 30 degrees to allow network silencing. Flies were collected in 6h intervals in the fifth day of constant darkness. Control flies (*tub-GAL80ts UAS-Kir*, black data points ± SEM) show cycling with peak expression around ZT0 in all subsets of neurons. Silencing the network causes the sLNvs to get arrhythmic (left panel). The LNds show no reduction in cycling amplitude and only a slight shift in protein cycling (middle panel). The DN1s (right panel) show a reduction of PER cycling amplitude.

**Supplement Figure 4.**
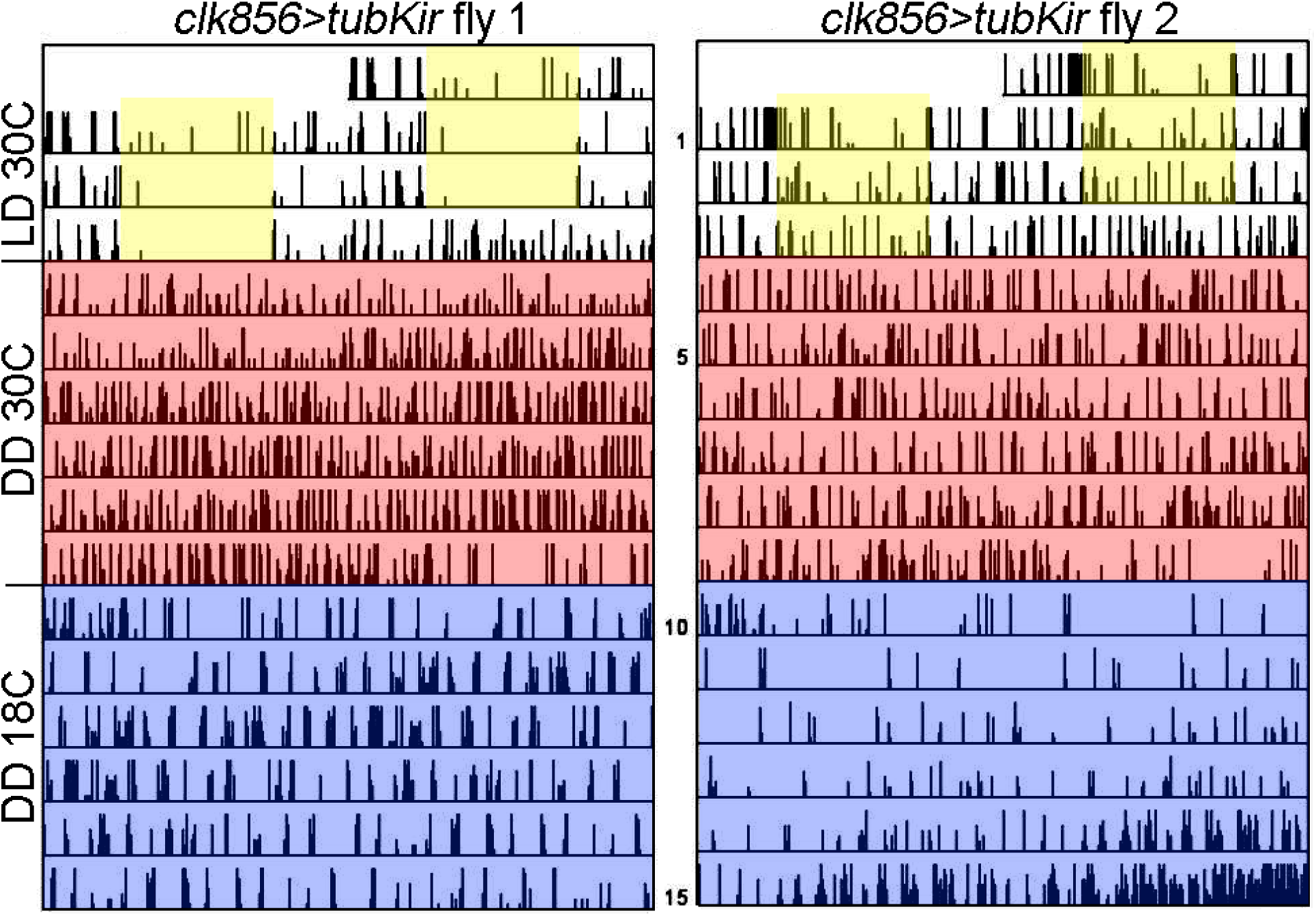
Stopping neuronal silencing after 6 days in DD does not re-establish rhythmic behavior. Actograms of individual flies recorded at 30 degrees for 3 days in LD 12:12 (indicated by yellow boxes). Flies were then transferred into constant darkness at 30 degrees (red area). Flies show no sign of rhythm due to the silencing of the network. We lowered the temperature to 18 degrees (blue area) to stop neuronal silencing. None of the flies re-established rhythmic behavior.

**Supplement Figure 5.**
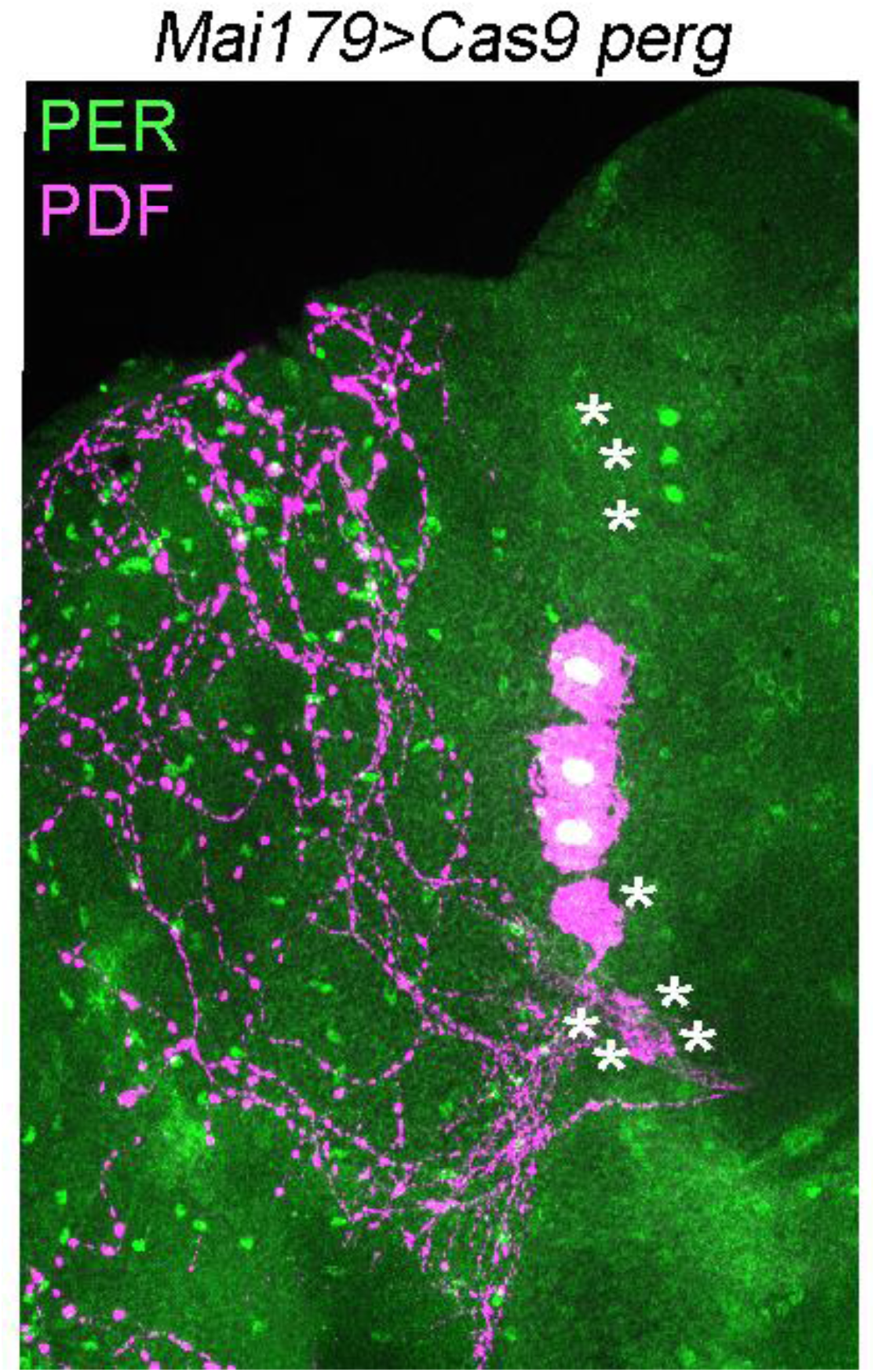
Immunolabeling of cell-specific per KO using *Mai179*-GAL4. *Mai179*-GAL 4 is expressed in 3 out of 6 LNds, the sLNvs and shows weakly and variable expression in DN1 and lLNv neurons. We performed PER (green) and PDF (magenta) in *Mai179*>*Cas9 per-g* flies and found that 3 LNds were PER+, the sLNvs were PER- and one of the lLNvs was PER-. This nicely reproduces the expression pattern mentioned above. Asterix represents successful KO.

**Supplement Figure 6.**
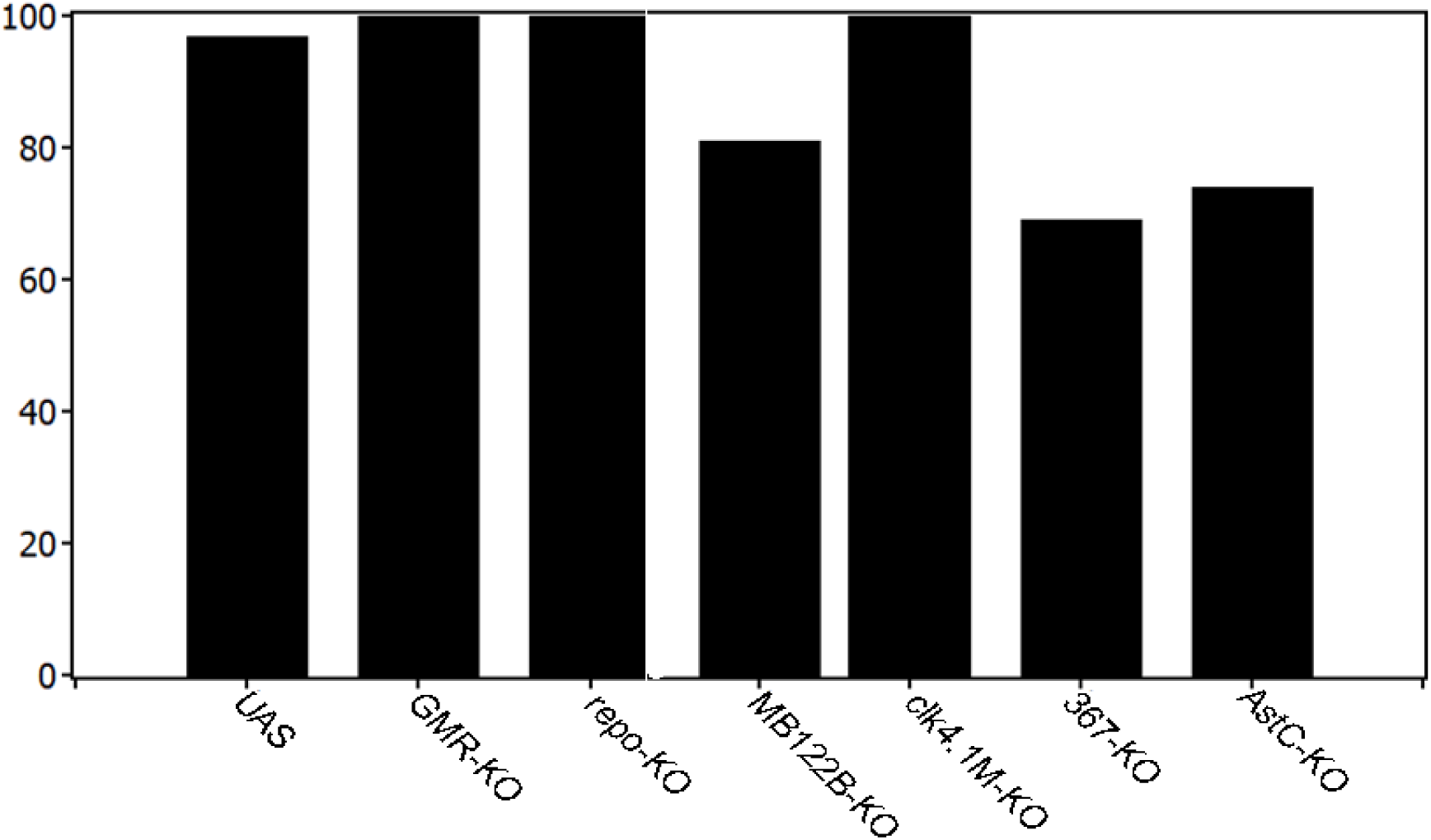
PER knockout in non-lateral neuron clusters does not affect rhythmicity levels. All investigated genotypes showed high levels of rhythmicity, suggesting that the LNds and sLNvs are the key players in determining rhythmicity.

## Notes

The authors declare no competing interests.

